# SHIP-1 differentially regulates IgE-induced IL-10 and antiviral responses in human monocytes

**DOI:** 10.1101/2024.02.07.579109

**Authors:** Siva Kumar Solleti, Bailey E. Matthews, Regina K. Rowe

**Author notes:** Corresponding author: Regina K. Rowe, MD PhD, University of Rochester Medical Center 601 Elmwood Ave, Box 690, Rochester, NY 14642.

## Abstract

IgE-mediated stimulation of monocytes regulates multiple cellular functions including cellular maturation, cytokine release, antiviral responses, and T cell priming and differentiation. The high affinity IgE receptor, FcεRI, is closely linked to serum IgE levels and atopic disease. The signaling molecules which regulate effector functions of this receptor have been well studied in mast cells and basophils, however, less is known about the signaling components, regulatory molecules, and mechanisms downstream of receptor activation in monocytes. This study sought to identify regulators of IgE-mediated cytokine release in human monocytes. SHIP-1 was identified as a negative regulator of IgE-induced IL-10 production. It was also determined that IgE-mediated stimulation and SHIP-1 inhibition decreased antiviral IP-10 production after liposomal poly(I:C) stimulation, indicating differential regulation by SHIP-1 in IgE-driven and antiviral response pathways. Both SHIP-1 and NF-κB were activated following IgE-mediated stimulation of primary monocytes, and NF-κB activation was related to both SHIP-1 and FcεRIα expression levels in monocytes. To our knowledge this is the first study to identify a role for SHIP-1 in regulating IgE-driven responses and antiviral responses in human monocytes. Given the importance of monocytes in inflammation and immune responses, a better understanding of the signaling and regulatory mechanisms downstream of FcεRI receptor could lead to new therapeutic targets in allergic disease.

## Introduction

Monocytes serve as powerful regulators of immune responses and are functionally impacted by the allergic environment [1, 2]. *In vivo*, monocytes are recruited to airways after allergen exposure [2–5] and recent research using next-generation sequencing approaches (*eg.* RNA sequencing) identified monocyte transcriptomic signatures that correlated with disease states in allergic asthma [6, 7]. Additionally, IgE-mediated stimulation of primary human monocytes *ex vivo* induces secretion of regulatory and pro-inflammatory cytokines, including IL-10, IL-6, and TNF-α [7–9], inhibits virus-driven Th1 differentiation [10, 11], and enhances Rhinovirus-induced Th2 development [11]. These findings support an important role for the IgE receptor in regulating monocyte functions in allergic disease, however, the cellular factors regulating receptor signaling and downstream gene regulation are not well understood.

The high affinity IgE receptor, FcεRI, is primarily expressed on immune cells including mast cells, basophils, dendritic cells (both plasmacytoid and myeloid DCs), and monocytes [12–14]. The receptor is either tetrameric, composed of an alpha (FcεRIα), beta, and two gamma chains in mast cells and basophils, or in antigen presenting cells (monocytes and dendritic cells) the receptor lacks the beta chain, existing as a trimer of only the alpha and two gamma subunits [13, 15]. This cell-type specific composition suggests differential signaling and gene regulation, which could influence cell-type specific effector functions. Much is known about the specific signaling and regulation of the FcεRI in mast cells and basophils. For example, IgE-mediated activation of FcεRI results in signal transduction via SH-2 domain receptor tyrosine kinases, such as Lyn and Spleen tyrosine kinase (Syk), terminating in NF-κB activation for downstream gene regulation [15]. Conversely, the SH-2 containing inositol 5’ polyphosphatase (SHIP-1) has been identified as an important negative regulator of receptor signaling [16–18]. While receptor signaling and regulatory molecules are thought to be conserved across immune cell types, the specific signaling components and regulatory mechanisms involved in IgE-mediated effector functions in human monocytes is less clear.

This study investigates the role of SHIP-1 in regulating IgE-mediated effector functions in human monocytes. We identified SHIP-1 as a negative regulator of IgE-mediated IL-10 production from primary human monocytes, as well a role for SHIP-1 in monocyte antiviral responses. To our knowledge this is the first study to identify a role for SHIP-1 in both IgE-driven and antiviral responses in human monocytes.

## Results

### IgE crosslinking activates NF-κB and SHIP-1 in human monocytes

Previous findings in mast cells and basophils demonstrate that IgE crosslinking activates multiple receptor tyrosine kinases, including Lyn and Syk, and the transcription factor NF-κB [19, 20], while the inositol 5’ polyphosphatase, SHIP-1, conversely, has been identified as a negative regulator of FcεRI signaling [16, 21]. While data has shown that both Syk and NF-κB are activated downstream of activation of the trimeric FcεRI receptor [22, 23], there are still knowledge gaps in the exact signaling components and how regulators such as SHIP-1 are involved in primary human monocytes. We wanted to determine whether IgE crosslinking of primary human monocytes induces activation of previously identified components of the FcεRI signaling pathway in mast cells and basophils, including Lyn, Syk, and NF-κB. We also sought to determine if SHIP-1, as a candidate negative regulator, is also activated by IgE crosslinking, and whether inhibition of SHIP-1 impacted activation of FcεRI-associated signaling molecules. To test this, we isolated primary human CD14+ monocytes from PBMCs obtained from discarded leukocyte fractions of blood bank donors (Table 1). Monocytes were pre-treated with a pharmacologic inhibitor of SHIP-1 (3AC, iSHIP) for 1 hour, followed by IgE crosslinking, media (carrier), or IgG isotype controls for 30 minutes. Western blotting was performed for phosphorylated forms of the kinases Lyn (p-Lyn, Tyr397) and Syk (p-SYK, Tyr525/526), as well as SHIP-1 (p-SHIP-1, Y1020), and the p65 subunit of NF-κB, (p-p65 (S536)) (Fig 1A-B). Monocytes after IgE crosslinking showed increased activation (*i.e.* phosphorylation) of all four of the FcεRI-associated signaling molecules (Fig. 1B). While IgE crosslinking reproducibly activated these molecules, SHIP-1 inhibition did not significantly affect either baseline or IgE-induced activation of these molecules (Fig. 1B).

**Figure 1.**
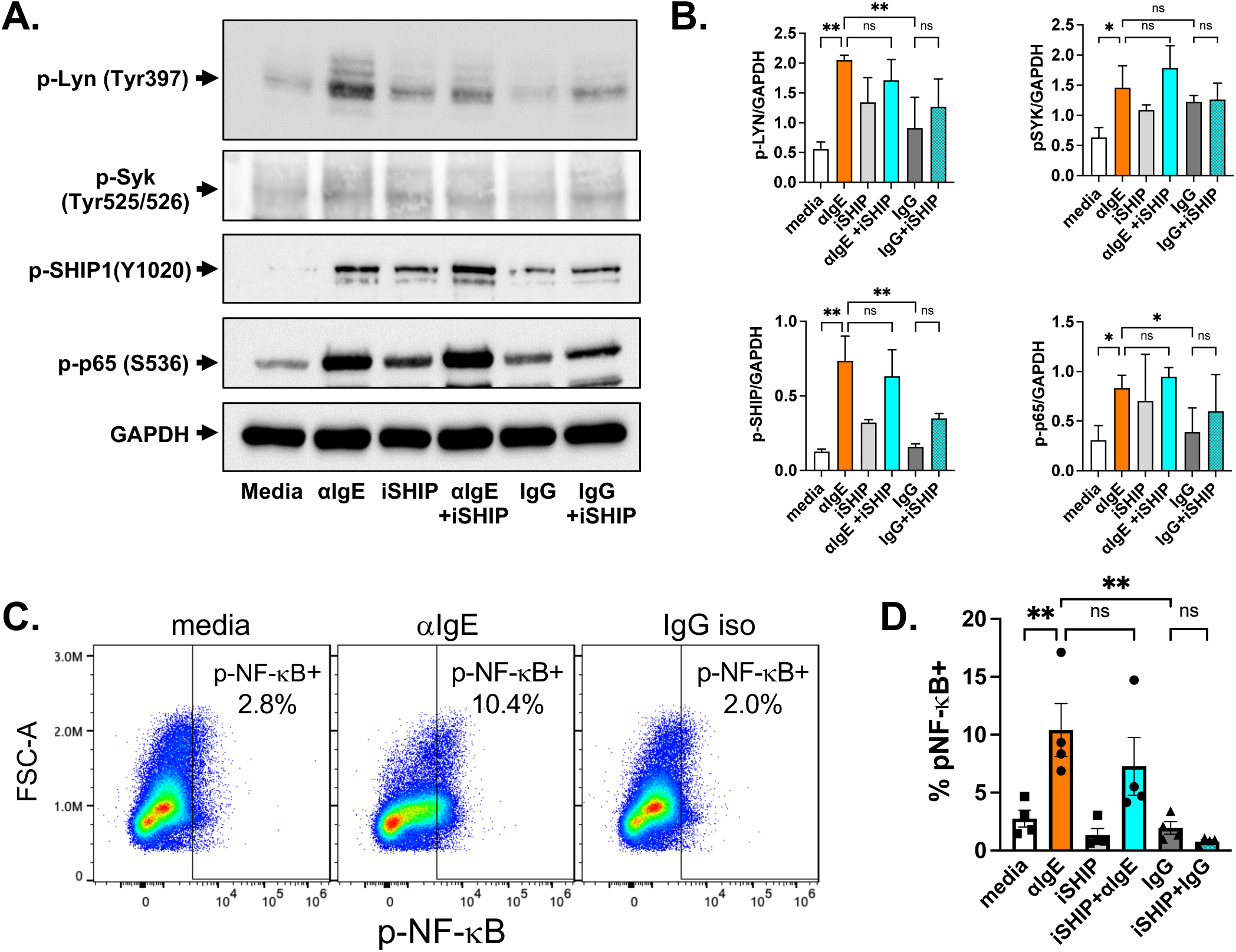
IgE-crosslinking activates NF-κB and SHIP-1 in monocytes. Primary human monocytes were pre-treated with SHIP-1 inhibitor, 3AC (iSHIP), for 1 hour then IgE crosslinking was performed (αIgE) along with media alone and IgG isotype controls for 30 minutes. (A & B) Whole cell lysates were analyzed by western blot analysis for phosphorylated forms of signaling molecules: p-Lyn, p-Syk, p-SHIP-1, and the p65 subunit of NF-kB (p-p65). GAPDH was used as a loading control for quantification. N=3 biological replicates were performed and mean with SEM are shown in (B). p values were obtained by one-way ANOVA with post-hoc comparisons. (C & D) Flow cytometry analysis was performed for phosphorylated p65 NF-κB (p-NF-κB). (C) Flow cytometry gating strategy for p-NF-κB+ cells in purified monocytes. Flow plots shown are a concatenation of 4 biological replicates to demonstrate the range of cellular data. (C & D) p-NF-κB positive cells were measured across the various treatment conditions; the mean with SEM shown of 4 biological replicates are shown in (D). p values obtained by ANOVA with Fishers LSD test post-hoc comparisons; *p<0.05 and **p<0.01.

**Table 1:**
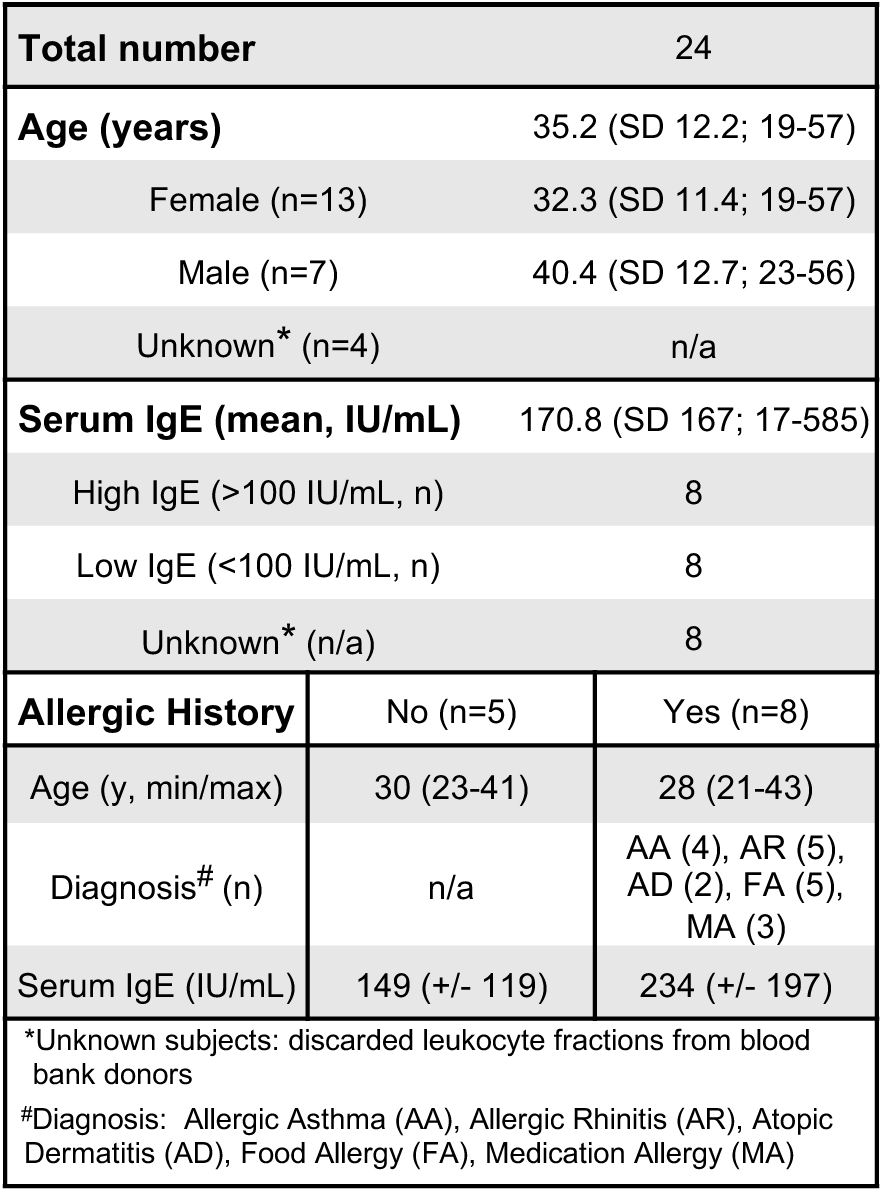
Subject Demographics.

### IgE-induced NF-κB activation is associated with SHIP-1 and FcεRIα expression levels

It has been shown that a subset of monocytes has higher expression of FcεRI, which is related to serum IgE levels [24, 25]. We hypothesized that IgE-induced NF-κB activation may be confined to a fraction of monocytes and differentially regulated by SHIP-1, which may not be detectable by bulk western blot analysis. To evaluate this on a single cell level, purified monocytes were pretreated with SHIP-1 inhibitor followed by IgE crosslinking for 30 minutes and evaluated by flow cytometry for phosphorylated NF-kB (p-p65) (Fig. 1C-D). Similar to western blot analysis, IgE crosslinking induced phosphorylation of NF-κB p65 as compared to media alone or IgG isotype control (p<0.01) with no effect of SHIP-1 inhibition (Fig. 1C-D).

Co-staining for total cellular SHIP-1, FcεRIα, and MHC II (HLA-DR) was also performed (Fig. 2) to determine if cells with activated (phosphorylated) NF-κB (p-NF-κB+) were associated with a specific cellular phenotype. We first compared expression of MHC II (HLA-DR), FcεRIα, and SHIP-1 in the p-NF-κB+ population with the total monocyte population (parent gate, MHC II+) (Fig. 2A-B). The p-NF-κB+ population had significantly higher expression of all three markers as compared to the total monocyte population (Fig. 2B). Similarly, we noted that there were two populations of monocytes with regards to SHIP-1 expression levels (Fig. 2C; SHIP1 hi, SHIP1 low). The SHIP-1 high population had higher MHC II expression as compared to low SHIP-1 expressing cells (p<0.01) in both media and IgE crosslinking conditions (Fig. 2D). The SHIP-1 high population also had an increased percentage expressing FcεRIα in media alone conditions (p<0.05) (Fig. 2D). While there was a trend in the IgE-crosslinking condition (Fig. 2D), this did not reach significance, likely due to potential downregulation of FcεRIα after IgE crosslinking [26]. Finally, the percentage of cells with p-NF-κB activation and amount of p-NF-κB were also higher in the SHIP-1 high population (Fig. 2E-F). These data indicate that the subset of monocytes activated by IgE crosslinking have higher expression of the high affinity IgE-receptor (FcεRI), antigen presenting molecules (MHC II), and SHIP-1.

**Figure 2.**
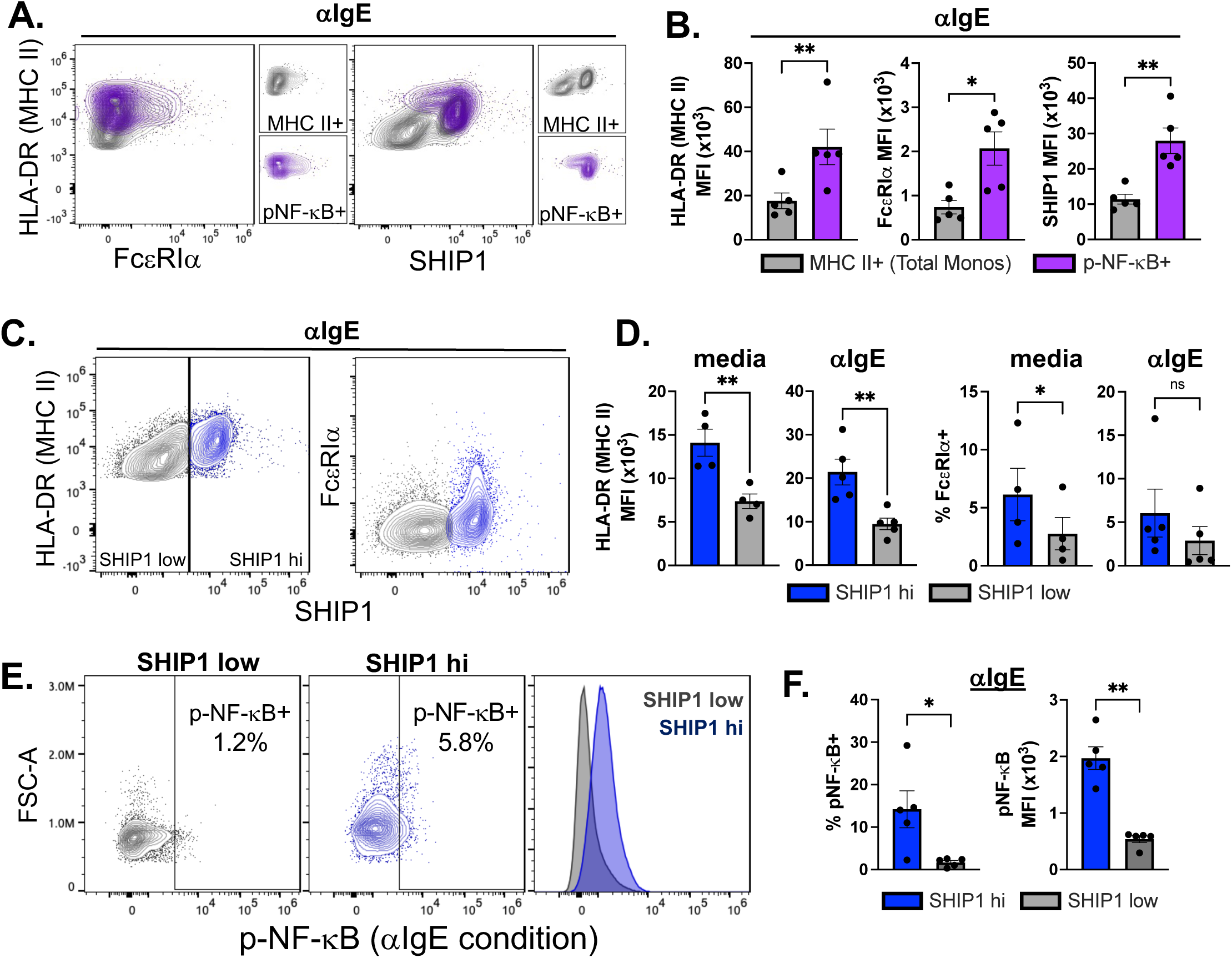
IgE-mediated activation of NF-κB is related to MHC II, SHIP-1, and FcεRIα expression. Primary human monocytes were pre-treated with SHIP-1 inhibitor, 3AC (iSHIP), for 1 hour then IgE crosslinking was performed (αIgE) along with media alone and IgG isotype controls for 30 minutes. Flow cytometry analysis was performed for phosphorylated p65 NF-κB (p-NF-κB), and total MHC II (HLA-DR), FcεRIα, and SHIP-1. Flow plots are a concatenation of all biological replicates to demonstrate the range of cellular data. (A & B) Under IgE crosslinking conditions (αIgE), MHC II, FcεRIα, and SHIP-1 levels were compared in the p-NF-κB positive population (purple) as compared to total monocytes (grey). (C & D) MHC II and FcεRIα were analyzed in the SHIP-1 high (blue) and low (grey) populations in media alone and IgE crosslinking conditions. (E & F) p-NF-κB levels were compared in SHIP-1 high and low cell populations. N=5; p values were obtained by paired two-tailed t test; *p<0.05 and **p<0.01.

### SHIP-1 is involved in IgE-mediated IL-10 production

IgE crosslinking of monocytes induces secretion of various pro-inflammatory and regulatory cytokines, including IL-6, IL-1β, TNF-α, IL-23, and IL-10 [8–11], but how IgE-activated signaling pathways regulate production of these cytokines is not known. In mast cells and basophils, both Syk and SHIP-1 have been shown to regulate IgE-induced cellular effector functions, including cytokine release [17, 19, 27, 28]. Given that IgE crosslinking of monocytes similarly resulted in activation of these molecules, we investigated the role of SHIP-1 and Syk in IgE-driven IL-10 and IL-6 production. Primary monocytes were isolated from PBMCs from adults with and without a history of allergy recruited from the local community, as well as unknown subjects, via discarded leukocyte fractions from blood banks (Table 1). Allergic history was obtained via self-report on a medical history questionnaire, and serum was collected for IgE levels (Table 1). Monocytes were pre-treated with inhibitors of either SHIP-1 (3AC, iSHIP) or Syk (R406, iSyk) followed by IgE crosslinking, and measurement of IL-10 and IL-6 levels at 24 hours post-crosslinking (Figure 3). Compared to media alone or IgG isotype control, IgE crosslinking induced significant secretion of IL-10 and IL-6 (Fig. 3). Inhibition of SHIP-1 resulted in a significant enhancement of IL-10 levels (p<0.01) (Fig.3A). Syk inhibition, however, did not have any significant effect on IgE-mediated IL-10 (Fig. 3B). SHIP-1 inhibition did trend towards decreasing IL-6 secretion (p=0.08) (Fig. 3C) with Syk inhibition slightly increasing its secretion (p=0.08) (Fig. 3D). The effects on IL-6 did not reach significance, suggesting that the role of SHIP1 is more specific to IgE-induced IL-10. We found no difference in IL-10 or IL-6 levels with or without SHIP inhibition based on serum IgE levels or allergic history (data not shown), which may suggest conservation in SHIP-1 mediated mechanisms in human monocytes regardless of atopic history. These results indicate that SHIP-1 negatively regulates IgE-induced IL-10 secretion from human monocytes.

**Figure 3.**
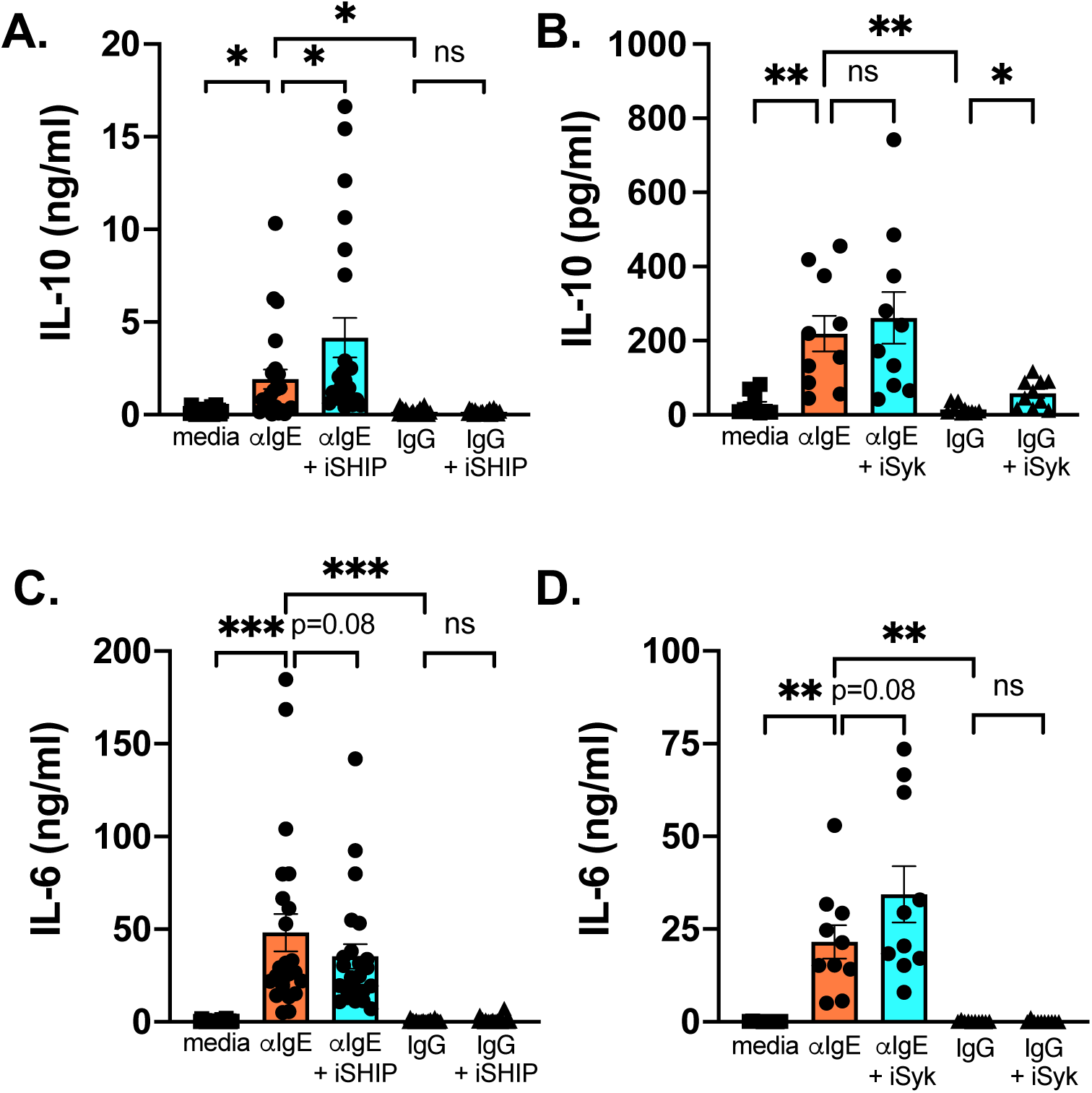
SHIP-1 inhibition enhances IgE-induced IL-10 production. Primary human monocytes were purified and treated with inhibitors to (A & C) SHIP-1 (3AC, iSHIP, n=23) or (B & D) Syk (R406, iSyk, n=10) for 1 hour followed by IgE crosslinking (αIgE) or IgG isotype or media (carrier) alone controls. Supernatants were harvested at 24 hours and (A & B) IL-10 and (C & D) IL-6 measured by ELISA. Bar graphs show mean with SEM with individual subject data points overlaid. p values obtained by ANOVA with mixed effects analysis and post-hoc comparisons; *p<0.05, **p<0.01, and ***p<0.001.

### SHIP-1 regulates both IgE-mediated and antiviral responses in human monocytes

IgE-mediated stimulation is known to inhibit monocyte antiviral responses to both influenza and rhinovirus [10, 11]. SHIP-1 has separately been shown to be involved in regulation of innate immune pathways [29]. However, SHIP-1 involvement in antiviral responses in human monocytes - and any role in IgE-mediated inhibition of antiviral responses - is not defined. In lieu of virus exposure, which can activate multiple antiviral response pathways, we activated the cytosolic RNA sensing pathway by using liposomal poly(I:C) treatment. We first evaluated the impact of IgE crosslinking on monocyte secretion of the early antiviral response protein, IP-10 (CXCL-10), in response to liposomal poly(I:C) stimulation (Fig. 4). Primary human monocytes secreted large amounts of IP-10 in response to liposomal poly(I:C) (p<0.01), which was significantly inhibited by IgE crosslinking (p<0.01), with no effect of IgG isotype control treatment (Fig. 4A). Next, we investigated the role of SHIP-1 on liposomal poly(I:C)-induced IP-10 secretion (Fig. 4B). SHIP-1 inhibition alone significantly inhibited liposomal poly(I:C) induced IP-10 production (p<0.05), and the combination of IgE crosslinking and SHIP-1 inhibition synergistically decreased IP-10 release as compared to either SHIP-1 inhibition (p<0.01) or IgE crosslinking alone (Fig. 4B). Since IgE-induced IL-10 has previously been shown to regulate antiviral responses in monocytes [11], we measured secretion of IL-10 (Fig. 4C) in the setting of liposomal poly(I:C). Similar to previous studies using live RNA virus exposure [10, 11], IL-10 secretion was specific to IgE crosslinking, as liposomal poly(I:C) did not trigger significant IL-10 release (Fig. 4C). However, liposomal poly(I:C) treatment in combination with IgE crosslinking significantly enhanced IL-10 release as compared to IgE-crosslinking alone (p<0.05) (Fig. 4C). We performed correlation analysis of IP-10 and IL-10 levels and found that in the setting of liposomal poly(I:C) treatment, IgE-induced IL-10 negatively correlated with the secretion of IP-10 (R^2^=0.309; p=0.039) (Fig. 4D). These results indicate that both IgE crosslinking and SHIP-1 regulate production of the early antiviral factor, IP-10, in response to liposomal poly(I:C) treatment in human monocytes, and IgE-induced IL-10 may play a role in IgE-induced suppression of antiviral IP-10.

**Figure 4.**
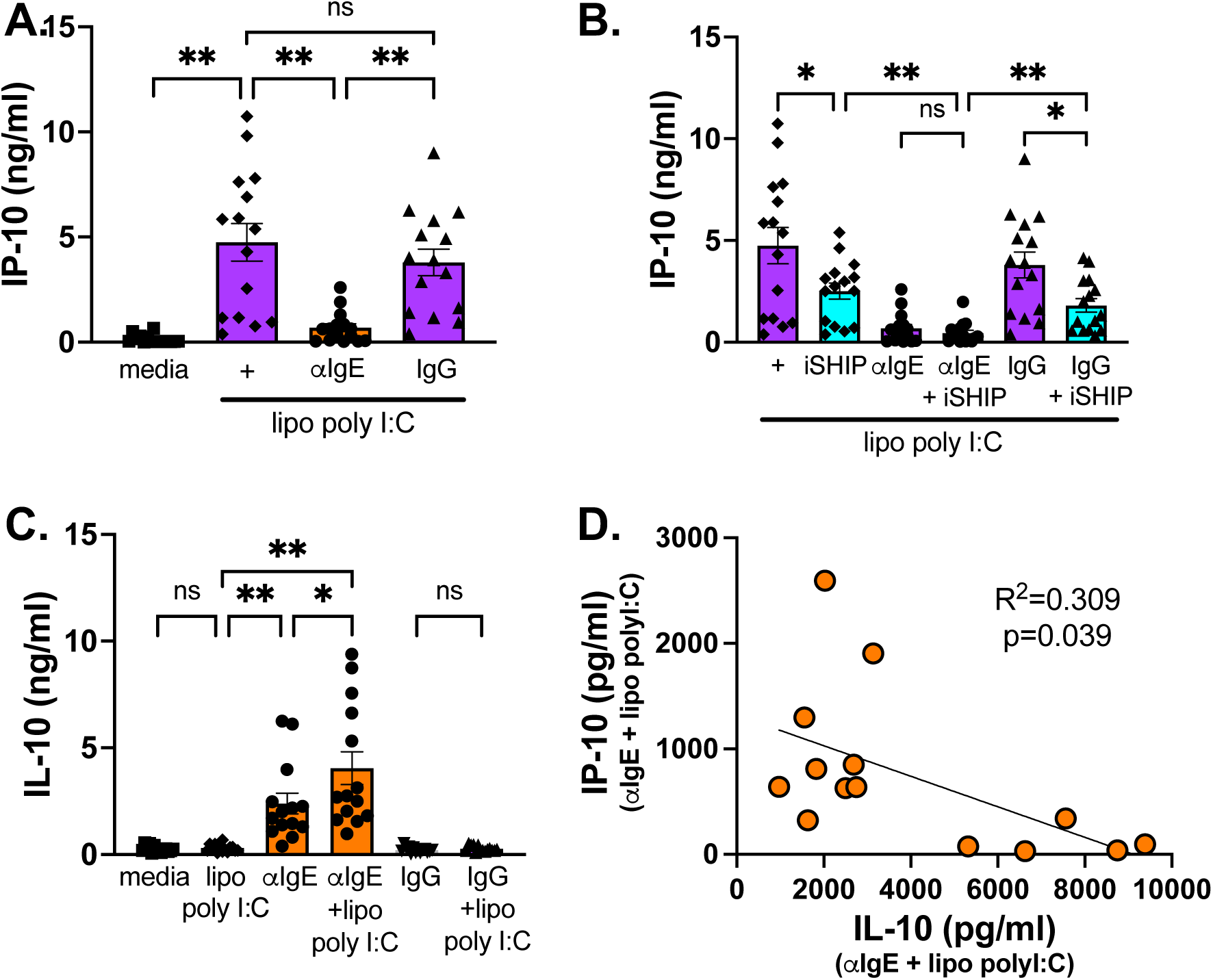
IgE-mediated stimulation and SHIP-1 regulate antiviral IP-10 in human monocytes. Primary human monocytes were pre-treated with or without SHIP-1 inhibitor, 3AC (iSHIP), for 1 hour then followed by IgE-crosslinking (with media or IgG isotype controls) for 1 hour, and subsequent stimulation with liposomal poly(I:C). Supernatants were harvested at 24 hours and analyzed for IP-10 and IL-10 release. (A & B) IP-10 production after liposomal poly(I:C) treatment with and without IgE crosslinking or IgG isotype control and (B) combined with SHIP inhibition. (C) IL-10 production after IgE crosslinking and liposomal poly(I:C) treatment. (D) Linear regression analysis of IL-10 versus IP-10 in the αIgE + liposomal polyI:C condition. Pearson coefficient (R^2^) and p value noted. Mean with SEM and individual subjects shown for n=15 biological replicates. ANOVA with mixed effects analysis and post-hoc comparison was performed; *p<0.05, **p<0.01 and “ns” p>0.05.

## Discussion

IgE has a critical role in the development and progression of atopic diseases, and signaling through the high affinity IgE receptor, FcεRI, is a key mediator of IgE-driven cellular effector functions. In this study, we sought to identify regulators of IgE-mediated cytokine release in human monocytes. We found a role for SHIP-1 as a regulator of both IgE-mediated IL-10 production and antiviral responses in primary human monocytes. Our model in Figure 5 proposes that SHIP-1 functions differentially across these pathways – as a negative regulator of IgE-induced IL-10 but potentially a positive regulator of antiviral IP-10 production.

**Figure 5.**
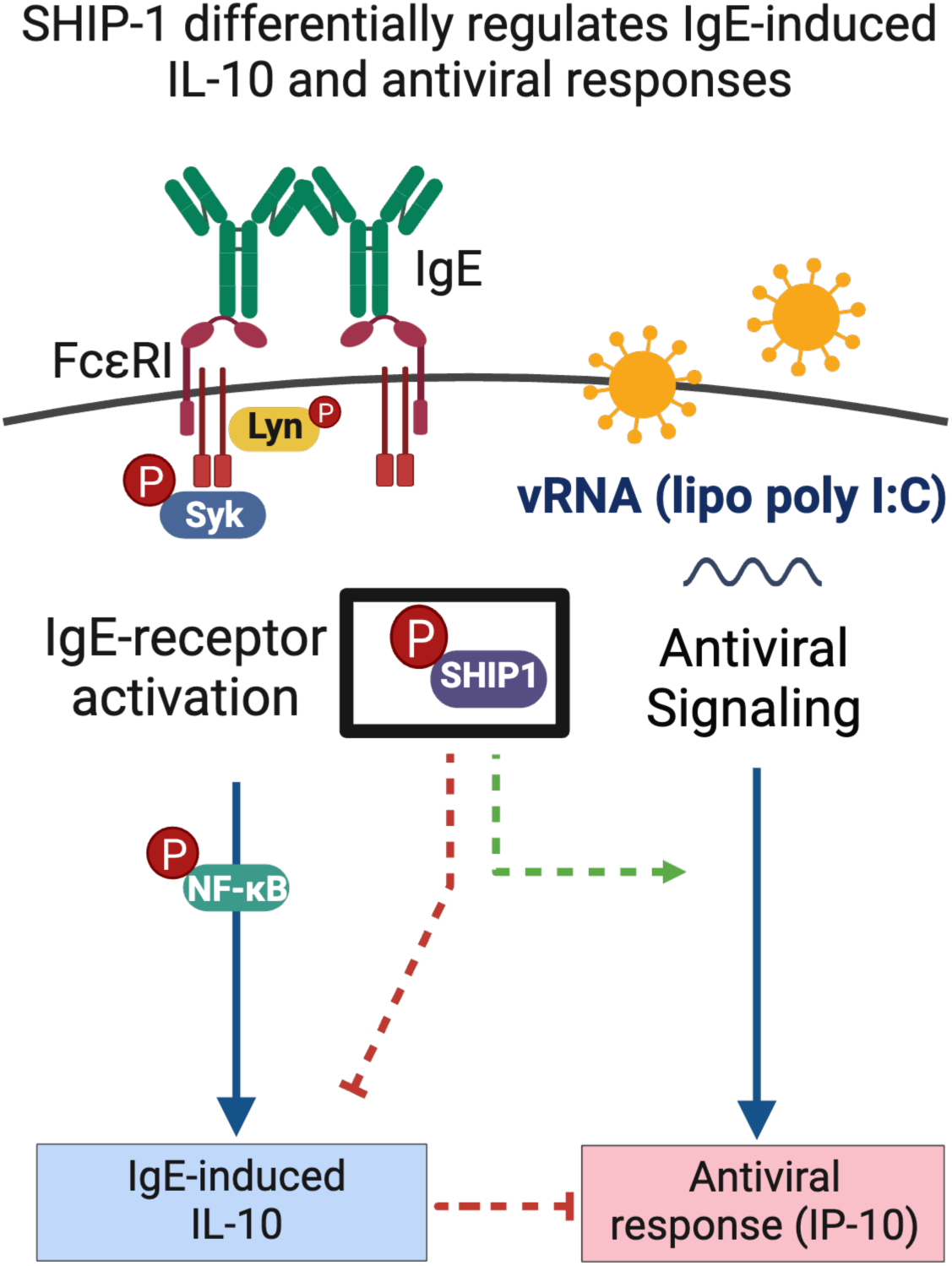
Model of SHIP-1-driven regulation of IgE receptor signaling and antiviral responses in monocytes. IgE-mediated activation of FcεRI leads to activation of downstream signaling molecules including receptor tyrosine kinases Lyn and Syk, and transcription factor, NF-κB via phosphorylation. SHIP-1 is also activated, serving as a negative regulator of IgE-induced IL-10. Simultaneously, IgE-mediated stimulation and SHIP-1 have differential effects on the regulation of liposomal poly(I:C) induced antiviral responses (IP-10 release).

While, FcεRI-mediated signaling is well described in mast cells and basophils, the signaling intermediates and regulators are not fully elucidated in human monocytes. In mast cells, FcεRI activation results in activation of receptor tyrosine kinases, Fyn and Lyn, followed by phosphorylation of ITAMs within the FcεRI β and γ subunits. This leads to docking of Syk to ITAMs and downstream cellular activation [19]. FcεRI-mediated NF-κB and Syk activation have been demonstrated previously in monocytes [23] and monocytic cell lines [30], respectively. We confirmed that IgE-mediated activation does trigger phosphorylation of Lyn, Syk, and NF-κB, consistent with previous findings in basophils and mast cells, indicating conservation of receptor signaling components. However, Syk inhibition did not block IgE-induced effector functions (IL-6 or IL-10 production) in our system, suggesting Syk is not required, and redundancy may exist in this signaling pathway in monocytes. Future work will include identifying the roles of other upstream receptor tyrosine kinases in regulating IgE-induced cytokine release.

SHIP-1 has been shown to be a critical negative regulator of FcεRI receptor activation [21]. IgE-induced activation of SHIP-1 in mast cells and basophils, serves to negatively regulate IgE-receptor signaling and effector functions [20, 21, 27]. Similarly, we found that SHIP-1 was phosphorylated after IgE crosslinking of human monocytes, indicating that this regulatory pathway may be conserved. Supporting its role as a negative regulator, pharmacologic inhibition of SHIP-1 resulted in increased IgE-induced IL-10 production (Fig. 3A). While phosphorylation of SHIP-1 at Y1020 is a marker for activity, phosphorylation at this site does not directly impact phosphatase activity [31] and pharmacologic inhibition with 3AC does not alter phosphorylation. Consistent with this, we detected IgE-induced phosphorylation in the setting of SHIP inhibition

(Fig. 1A & B), indicating that phosphatase activity was an important regulatory step in IgE-induced IL-10 production. Interestingly, SHIP-1 inhibition did not have the same effects on IgE-induced IL-6 production as has been shown in mast cells [27]. There was a trend to decreased IL-6, suggesting differential regulation of IgE-induced cytokines (Fig. 3C). We hypothesize that the decrease in IL-6 is likely related to increased IgE-induced IL-10 production in the context of SHIP-1 inhibition, as IL-10 is a known negative regulator of IgE-induced IL-6 [8]. Additional studies are necessary to identify how IgE-driven cytokines, such as IL-10 and IL-6, are differentially induced and the mechanisms by which SHIP-1 regulates IgE-mediated IL-10, selectively.

SHIP-1 has multiple described roles within the cell [29], thus may be functioning at multiple stages of IgE-mediated effects. It has been shown to act as a negative regulator of NF-κB activity in macrophages [32] and impact IgE-driven activation of NF-κB in mast cells [27]. We showed that IgE-crosslinking activates NF-κB in human monocytes, consistent with previous reports (Fig. 1 & 2). SHIP-1 inhibition did not impact IgE-driven phosphorylation of NF-κB, Lyn, or Syk, supporting alternative mechanisms downstream of IgE-receptor signaling and NF-κB activation by which SHIP-1 is functioning. Our flow cytometry results indicated differential activation across monocytes subsets, as monocytes with high levels of NF-κB activation were higher for both SHIP-1 and FcεRIα levels. Similarly, SHIP-1 high monocytes also had a higher percentage of phosphorylated NF-κB and higher total FcεRIα expression (Fig. 2). These data suggest that monocytes higher in IgE receptor expression may have differential activation by IgE crosslinking and subsequent SHIP-1 regulation as compared to the total monocyte population. While we did not find an effect of serum IgE levels or allergic history on enhanced IL-10 production after SHIP-1 inhibition, our flow cytometry data support a relationship between SHIP-1, FcεRI, and IgE-mediated activation in human monocytes. Future studies addressing co-regulation of these pathways and molecules - at the single cell level - in a larger cohort of well characterized atopic subjects, could elucidate IgE-driven differences in SHIP-1-regulated pathways specific to monocyte subsets and atopic disease.

In both monocytes and dendritic cells, IgE-crosslinking inhibits antiviral responses, including inhibition of virus-induced type I IFN [10, 11, 33, 34], cellular maturation [10, 33], and virus-driven T cell differentiation [10, 11]. Links between SHIP-1 and regulation of antiviral signaling pathways have been shown. For example, SHIP-1 deficient murine macrophages have constitutive activation of TBK1 (Tank-binding kinase 1), a kinase critical for antiviral signaling [35], and loss of SHIP-1 in mice alters LPS-induced IFNβ production [36]. We demonstrated that IgE-mediated stimulation of monocytes inhibits IP-10 (CXCL10) production after activation of the cytosolic RNA sensing pathways by liposomal poly(I:C). SHIP-1 inhibition did not reverse IgE-mediated inhibition, but instead suppressed liposomal poly(I:C) induced IP-10. IP-10 was further inhibited synergistically by the combination of SHIP-1 inhibition and IgE crosslinking (Fig. 4). This indicates that while IgE crosslinking negatively regulates antiviral responses, SHIP-1 is acting as a positive regulator of liposomal poly(I:C) induced IP-10 (Fig. 5).

IL-10 is a negative regulator of many pro-inflammatory cytokines, including IP-10 [37], as well as itself in the setting of IgE-crosslinking [8]. Interestingly, we observed that the combination of IgE crosslinking and liposomal poly(I:C) treatment led to an enhancement in IgE-induced IL-10 (Fig. 4). While this was not previously observed in the context of live virus exposure [10, 11], a similar result was reported in myeloid DCs where IFNα treatment enhanced IgE-driven IL-10 production [38]. We hypothesize that the synergy in decreasing IP-10 between IgE crosslinking and SHIP-1 inhibition could be related to enhanced IL-10 production, given the observed negative correlation between IL-10 levels and IP-10 suppression in the setting of IgE crosslinking (Fig. 4 & 5). Our data supports cross-talk between the cytosolic RNA antiviral sensing and IgE-mediated pathways. This co-regulation of IL-10 and antiviral responses has downstream implications for allergic inflammation. As shown previously, IgE-induced suppression of virus-induced IFNα by IL-10 in monocytes promotes virus-driven Th2 differentiation *ex vivo*, which could further enhance allergic inflammatory responses [11].

To our knowledge this is the first study in primary human monocytes demonstrating IgE-induced activation of SHIP-1 and a role for this molecule as a negative regulator of IgE-mediated IL-10 production (Fig. 5). We also identified involvement of SHIP-1 in monocyte antiviral IP-10 production, including co-regulation by SHIP-1 with IgE-mediated effects in antiviral responses. Thus, our data support a role for SHIP-1 acting as both a positive and negative regulator in human monocytes with potential cross-talk between FcεRI and antiviral pathways (Fig. 5). Future work is needed to fully elucidate the molecular mechanisms of how SHIP-1 is regulating IgE-mediated and antiviral pathways in atopic disease.

## Materials and methods

### Human Subjects

Primary human immune cells were isolated from 2 sources: leukocyte-enriched blood samples from unknown subjects acquired from blood banks and peripheral blood samples from healthy and allergic subjects (Table 1). Unknown subjects were deemed healthy enough for routine blood donation by the blood bank, with limited demographic information (age and sex) available. In a select subset of these unknown subjects, we were able to obtain serum to evaluate IgE levels, otherwise no clinical information (*i.e.* atopic status or allergic history) was obtained. For all known subjects, written informed consent and assent were obtained. Known subjects were recruited from the local Rochester community and completed a medical and allergy history questionnaire. Peripheral blood was obtained to isolate PBMCs and serum was collected to measure IgE levels. Studies were approved by the University of Rochester Research Studies Review Board (RSRB) (Study #00006081).

### Serum IgE levels

Serum was isolated from 2-5mL of peripheral blood by serum separator tubes by spinning at 1200 x g for 10 minutes. Serum was harvested and frozen in 0.5mL aliquots at −80°C until analysis. To measure serum IgE levels, frozen serum was sent to the University of Rochester Medical Center Clinical Microbiology Laboratory and IgE levels were measured via a clinically validated test. Levels were reported in international units (IU) per mL. Subjects were divided into high IgE (>100IU/mL) or low IgE (<100IU/mL) groups, and by allergic history as shown in Table 1 for data analysis purposes.

### Reagents and Media

Phosphate Buffered Saline (PBS) with fetal bovine serum (FBS) and EDTA (complete PBS, cPBS) and complete Roswell Park Memorial Institute Medium 1640 (cRPMI) were prepared as previously described [8, 10, 11]. All medias were purchased from Gibco unless otherwise stated and FBS was purchased from Biowest (Riverside, MO). Goat anti-human IgE (αIgE) was purchased from Bethyl Laboratories (Montgomery, TX) and goat whole IgG isotype control antibody (IgG) from either Bethyl Laboratories or Jackson ImmunoResearch (Westgrove, PA). Goat anti-human IgE was chosen to perform IgE crosslinking for these studies to facilitate western blot analysis using rabbit anti-human antibodies to the various signaling molecules. Poly(I:C) (low molecular weight) LMW was purchased from InvivoGen, San Diego, CA, and Lipofectamine® 2000 was procured from Thermo Fisher Scientific, Waltham, MA. All the western blot buffers, Nitrocellulose Membrane, 0.45 µm, goat anti-rabbit or anti-mouse IgG HRP conjugates were purchased from Bio-Rad laboratories (Bio-Rad laboratories Inc., Hercules, CA). Rabbit monoclonal antibodies against Phospho-SHIP-1 (Tyr1020), Phospho-Syk (Tyr525/526) (clone C87C1), Phospho-LYN (Tyr397)/LCK (Tyr394)/HCK (Tyr411)/BLK (Tyr389) (clone E5L3D) and Phospho-NF-κB p65 (Ser536) (clone 93H1) were from Cell Signaling Technology, Inc. (Danvers, MA). Mouse monoclonal anti-GAPDH (clone 6C5) was purchased from Santa Cruz Biotechnology, Inc. (Dallas, TX). SuperSignal™ West Femto Maximum Sensitivity Substrate and Halt™ Protease and Phosphatase Inhibitor Cocktail were purchased from Thermo Scientific (Rockford, IL). Chemical inhibitors, 3 a-aminocholestane (3AC) (EMD Millipore, Burlington, MA) and R406 (InvivoGen, San Diego, CA) were dissolved in Ethanol (Koptec, PA) and DMSO (EMD Millipore, Burlington, MA), respectively. Resazurin, Sodium Salt was purchased from Thermo Scientific (Rockford, IL), and Ficoll-Paque™ PLUS from Cytiva (Uppsala, Sweden).

### Purification of Immune Cells from Blood

PBMCs were isolated from either discarded leukocyte fractions from blood bank donors or peripheral blood from known recruited subjects via Ficoll gradient centrifugation. PBMCs were then preserved frozen at 1×10^7^ cells per mL in freezing medium (10% DMSO, 50% FBS, 40% cRPMI). Cells were slow cooled at −80°C overnight in a Mr. Frosty^TM^ container (Thermo Scientific), then transferred to liquid nitrogen for long term storage. To isolate monocytes, PBMCs were quick thawed in a 37°C water bath for 2-3 minutes then washed once in cRPMI at 4 times the volume, pelleted and washed once more in cPBS. PBMCs were counted for viability with trypan blue (typically >90%) and resuspended at up to 5×10^7^ cells/mL in cPBS. Monocytes were purified by negative selection from PBMCs as previously [8, 10, 11] using the EasySep^TM^ Human Monocyte Enrichment kit (catalog #19059), per manufacturer’s instructions (STEMCELL Technologies, Vancouver). Purity (typically >85%) was assessed by flow cytometry with monocytes were identified as CD14+.

### Monocyte Culture Conditions

Monocytes were cultured in cRPMI at a concentration of 1 x 10^6^ cells/ml. As a surrogate for IgE-mediated allergic stimulation, we utilized previously published methods to crosslink cell surface bound IgE using a goat anti-human IgE polyclonal antibody (αIgE; Bethyl Laboratories, Montgomery, TX) [34]. While all individuals have surface bound IgE on monocytes, the level is related to serum IgE concentrations and thus varies across subjects [39–41]. Even with this variation, we have previously shown that this method of direct crosslinking induces reproducible IgE-mediated phenotypes in monocytes such as cytokine release [8] and inhibition of influenza-induced Th1 differentiation [10]. To best simulate the *in vivo* environment, monocytes were cultured and treated immediately after purification and any pre-treatments with inhibitors to ensure adequate crosslinking of native surface bound IgE and minimize any downregulation of surface receptors. Cells were treated with the following: αIgE (10 μg/ml), goat IgG isotype control (10 μg/ml). The following inhibitors were used to inhibit Syk kinase (R406, 7.5 μM/ml) and SHIP-1 (3AC, 7.5 μM/ml). For both inhibitors, the experimental concentration was determined based on previously published studies [42, 43] and were tested for cellular toxicity using resazurin assays (data not shown). The concentrations chosen had minimal/no loss of cellular viability and were comparable to effective concentrations used in other published studies [42, 43]. When inhibitors were used, cells were treated for 1-2 hours with inhibitor or carrier (in media) control then followed by IgE crosslinking and other stimulation (*eg.* liposomal poly(I:C)). To activate antiviral cellular responses, cells were treated with liposomal poly(I:C) (low molecular weight poly(I:C)). Liposomal poly(I:C) complexes were made by mixing poly(I:C) with Lipofectamine 2000 at a ratio of 1µg of poly(I:C) to 1µl of Lipofectamine 2000 reagent in RPMI media. The mixture was incubated at room temperature for 30 minutes and added to monocyte cultures at a concentration of 1µg/mL (based on poly(I:C) concentration). Cultures were incubated at 37°C for indicated times. Cells and supernatants were harvested for further analysis.

### Flow Cytometry Analysis

For flow cytometry analysis of phosphorylated (activated) NF-κB, total SHIP-1, and total FcεRIα, monocytes were stained with the following antibodies (antigen, clone, and fluorochrome noted respectively): FcεRIα (AER-37 (CRA-1), BV421), anti-SHIP-1 (P1C1-A5, PE), human IgE (MHE-18, Alexa Fluor 647), HLA-DR (L243, APC-Cy7) (all from Biolegend); and phospho-p65 NF-κB (K10-895.12.50, PE-Cy7) (BD Biosciences, San Jose, CA). For flow cytometry staining, purified monocytes were pre-treated with SHIP-1 inhibitor (3AC) for 1 hour, then followed with anti-IgE crosslinking or IgG isotype and media (carrier) alone controls for 30 minutes. Reaction was terminated by direct fixation with 1.6% paraformaldehyde for 10 minutes. Cells were washed once with cPBS and resuspended in 2mL of ice-cold methanol. Cells were preserved at −20°C until analysis. For flow staining, cells were re-hydrated as follows: 2mL of cPBS was added directly to the 2mL of methanol fixed cells, mixed, and pelleted for 10 minutes at 800 x g, then washed one additional time in 2mL of cPBS. Cells were pelleted and resuspended in antibody mixture diluted in cPBS and stained for 1 hour at room temperature. Samples were washed and resuspended in cPBS, then were acquired on a Cytek Aurora spectral flow cytometer (Cytek Biosciences, Fremont, CA) and analyzed using FlowJo^TM^ v10.10 (BD Life Sciences).

### Cytokine Secretion Analysis

Supernatants were stored at −80°C until use. The following ELISA kits were used according to manufacturer recommendations: ELISA Max human ELISA for IL-6, IL-10, or CXCL10 (IP-10) (Biolegend, San Diego, CA), Absorbance was measured on SpectraMax iD3 microplate reader (Molecular Devices, San Jose, CA) per manufacturer’s instructions.

### Western blotting of IgE-induced signaling molecules

Purified monocytes were treated as above with SHIP-1 inhibitor and then IgE crosslinked for 30 minutes. Cells were pelleted, washed in ice cold PBS- and directly lysed in 3X Laemmli sample buffer (188 mM Tris-Cl (pH 6.8), 3% SDS, 30% glycerol, 0.01% bromophenol blue and 15% β-mercaptoethanol) with protease and phosphatase inhibitor cocktail. Samples were boiled for 10 min and clarified by centrifugation. Cell lysates were run on SDS-PAGE and transferred on to nitrocellulose membranes and incubated with either 5% (w/v) BSA or 5% (w/v) nonfat dry milk in Tris-buffered saline with Tween-20 (TBST, 10 mm Tris pH 8.0, 150 mm NaCl, and 0.05% Tween-20) for 1 hr at room temperature. The membranes were then incubated with appropriate primary and secondary antibodies and developed using ECL method as per the manufacturer’s instructions and described previously [44]. Membranes were imaged using the Bio-Rad ChemiDoc MP (Bio-Rad, Hercules, CA). The intensity of each band in Western blot was quantified by using ImageJ software (National Institutes of Health, Bethesda, MD, USA).

### Data Analysis and Statistics

The number of biological replicates for a given experiment are noted in the figure legends. When cell numbers allowed, technical replicates were performed for IgE-induced cytokine and ELISA assays. Data in bar graphs are presented as the mean with standard error, with individual biological replicates overlaid. Other data are presented as means ± standard error of the mean (SEM) or standard deviation (SD) as noted. In experiments containing 3 or more conditions and comparisons, ANOVA with mixed effects analysis followed by post hoc comparisons were performed using Fisher’s LSD test, or Holm-Sidak correction for multiple comparisons performed when appropriate. For experiments comparing 2 conditions, paired two-tailed t tests were performed with Welch’s correction. For continuous variable analysis, linear regression analysis with two-tailed t-test was performed and Pearson’s coefficients (R^2^) and p values determined. p<0.05 was considered significant, pertinent p values are denoted as follows: *p<0.05, **p<0.01, and ***p<0.001, or ns (non-significant) for p>0.05. All statistical analyses were performed using GraphPad Prism version 10.

## Acknowledgements

We thank the study subjects, without whom this study would not be possible. We want to thank the URMC Vaccine Treatment and Evaluation Unit (VTEU), Dominique Williams, and Michelle Heiman (Division of Pediatric Infectious Disease, URMC) for assistance with study subject recruitment and study visit completion. We acknowledge the staff of the University of Rochester Medical Center Flow Cytometry Core for assistance with flow cytometry analysis. Figure 5 was created with Biorender.

## Funding Sources

NIH NIAID, 5K08AI163380 (RKR) and the Department of Pediatrics, University of Rochester Medical Center.

## Conflict of Interest Disclosure

The authors declare no commercial or financial conflict of interest.

## Contributions

SKS: Experimental design, data acquisition, data analysis, conceptualization, manuscript preparation

BEM: Data acquisition, data analysis, manuscript preparation

RKR: Experimental design, data analysis, conceptualization, manuscript preparation, supervision

## Abbreviations

αIgE: Goat anti-human IgE antibody, IgE crosslinking condition
CXCL: C-X-C motif chemokine ligand
DC: dendritic cell
FcεRI: high-affinity IgE receptor
FcεRIα: alpha subunit of the high-affinity IgE receptor
IgE: immunoglobulin E
IgG: Goat whole IgG isotype control antibody
IL: interleukin
IFN: interferon
IP-10: IFN-gamma-inducible protein 10 (CXCL10)
lipo pI:C: liposomal complexed low molecular weight (LMW) poly(I:C)
PBMCs: peripheral blood mononuclear cells
SHIP-1: SH-2 containing inositol 5’ polyphosphatase-1
SYK: Spleen tyrosine kinase

## Notes

### Competing Interest Statement

The authors have declared no competing interest.

